# Variance in Calvin-Benson cycle intermediate levels between closely-related species in the tomato clade

**DOI:** 10.64898/2026.02.28.708697

**Authors:** Vittoria Clapero, Stéphanie Arrivault, Mark Stitt

## Abstract

Published studies have reported species-variance between profiles of Calvin-Benson cycle (CBC) intermediates, not only between C_4_ species and C_3_ species, but also within C_3_ species (Arrivault *et al*., 2019, Borghi *et al*. 2019). It was proposed that this variance reflects lineage-dependent changes in the balance between different reactions, or poising, of the CBC. These earlier studies investigated phylogenetically-unrelated C_3_ species. In the current study, CBC intermediates were profiled in five closely-related species from *Solanum* sect. *lycopersicon* subsect. *Lycopersicum*. The levels of individual CBC intermediates showed many significant differences. In a principal component analysis, whilst three species (*Solanum lycopersicum, Solanum cheesmaniae, Solanum neorickii*) overlapped, *Solanum pimpinellifolium* and especially *Solanum pennellii* grouped separately, and were at opposing ends of the distribution. When combined with published data, whilst the separation between Solanum species was retained, they formed a group that was separated from five other C_3_ species, as well as two C_4_ species. It is discussed that the observed variation in CBC metabolites profiles within *Solanum*, together with their separation from other C_3_ species, supports the idea that CBC evolution is shaped both by phylogenetic relatedness and lineage-specific adaptation.

**Highlight:** Variance of intermediate levels points to poising of the Calvin-Benson cycle varying between closely-related species in the tomato clade *Solanum* sect. *lycopersicon* subsect. *Lycopersicum*

## Introduction

Since the Calvin–Benson cycle (CBC) evolved in cyanobacteria ∼2 billion years ago (Rasmussen *et al*., 2008) there have been large changes in the conditions under which it operates. Over geological time, there has been a dramatic decline in atmospheric CO_2_ and increase in atmospheric O_2_, including in the last 30 million years, when parallel evolution of C_4_ photosynthesis in >60 lineages coincided with a decline of atmospheric CO_2_ from ∼1000 ppm to <300 ppm (Sage *et al*., 2012; Raven *et al*., 2017). Species that did not evolve C_4_ photosynthesis are termed C_3_ species, representing ∼90% of existing terrestrial plant species (Silvera *et al*., 2010; Sage, 2017), will also have been subject to massive selective pressure by falling CO_2_, as well as its subsequent rise. Pressure will also have been exerted by further environmental factors like light, temperature and water and nutrient availability (Raven *et al*., 2017).

There is substantial variation in photosynthetic rate between terrestrial C_3_ species (Evans, 1989; Wullschleger, 1993), including between phylogenetically-related species (Galmés *et al*., 2014; Kromdijk and McCormack, 2022) and within species (Driever *et al*., 2014; Flood 2019). Contributing factors include differing rates of electron transport and carboxylation (Wullschleger, 1993), leaf nitrogen content, nitrogen use efficiency (Field and Mooney, 1986; Evans, 1989; Hikosaka, 2010), stomatal responses (Lawson and Vialet-Chabrand, 2019) and investment strategies in short- and long-lived leaves (Wright *et al*., 2004; Donovan *et al*., 2011). At the level of individual steps, it is well-established that Rubisco kinetics vary, even between quite closely related species (Yeoh *et al*., 1980; Galmés *et al*., 2014*a*; Prins *et al*., 2016; Hermida-Carrera *et al*., 2017). C_3_ species differ in how Rubisco is regulated by low molecular weight inhibitors (Servaites *et al*., 1986; Moore *et al*., 1993; Charlet *et al*., 1997; Parry *et al*., 2008). Regulation by CP12, a small protein that interacts with NADP-glyceraldehyde-3-phosphate and phosphoribulokinase, also varies between species (Howard *et al*, 2011). Recent studies in crops have uncovered variance in Rubisco capacity (V_cmax_) (Acevedo-Siaca *et al*., 2020, 2021), heat tolerance of Rubisco activase (Scafero *et al*., 2019), steady state and transient rates of photosynthesis (Pignon *et al*., 2021), heat stability of photosystem I (Posch *et al*., 2022) and cold tolerance (Ortiz *et al*., 2017). Based on evolutionary modelling, Zhu *et al*. (2007) proposed that current-day terrestrial plants may not yet have fully rebalanced the CBC after the low-CO_2_ and low temperature conditions they experienced in recent ice ages.

In recent years, the detection and mapping of allelic variance has provided an important and unbiased approach to studying variation in photosynthesis (Adachi *et al*., 2019; Faralli and Lawson, 2020; Kromdijk and McCormack, 2022; Sharwood *et al*. 2022, Sakoda *et al*., 2022). Metabolite profiling provides another unbiased approach to detect inter-specific variation, specifically in CBC operation. The underlying assumption is that changes in the relative levels of pathway intermediates reflect changes in the balance between enzymatic steps, or poising, in the pathway. Changes in poising are detected, regardless of whether they are due to changes in enzyme abundance, enzyme properties or regulatory loops. Liquid chromatography linked to tandem mass spectrometry (LC-MS/MS) and stable isotope standards allows quantitative profiling of most CBC intermediates (Arrivault *et al*., 2009; 2015; 2019) A diverse set of C_3_ species including the dicots *Arabidopsis thaliana, Nicotiana tabacum* (tobacco), *Manihot esculenta* (cassava) and *Flaveria* spp. and the monocots *Oryza sativa* (rice) and *Triticum aestivum* (wheat) showed significantly differing levels of many CBC metabolites (Arrivault *et al*., 2019; Borghi *et al*., 2019; 2022; Stitt *et al*., 2021). When displayed using principal component (PC) analysis, the changes distinguished one C_3_ species from another. These results were obtained for plants grown in moderately limiting irradiance and non-stressful conditions, and metabolites were profiled in material harvested in growth conditions. Whilst environment influences intermediate levels, the cross-species variance was interpreted as involving the impact of genotype or at least a genotype *x* environment interaction. The contribution of genotype was underlined by the finding that *A. thaliana* and *O. sativa* showed consistently different levels of CBC intermediates across a range of harvest irradiances (Borghi *et al*., 2019). These observations led to the proposal that this variability arises from continual selection on the CBC, and that this occurs in a lineage-dependent manner with different lineages following different adaptation trajectories (Arrivault *et al*., 2019; Stitt *et al*., 2021).

In previous studies, species were chosen based on their role as model organisms or important crops and, phylogenetically, were widely separated. A set of *Flaveria* C_3_ species, C_3_-C_4_ intermediate species and C_4_ species was studied to gain insights into the evolution of C_4_ photosynthesis (Borghi *et al*., 2022), but did not provide information about variation between phylogenetically-related C_3_ species. The *Solanum* genera, with ∼1500 species, is one of the largest genera of angiosperms. It originated in tropical South America, but spread worldwide. The tomato clade *Solanum* sect. *lycopersicon* subsect. *Lycopersicum* includes the cultivated tomato *Solanum lycopersicum* and 12 wild species (Peralta *et al*., 2008; Knapp and Peralta, 2016, Supplementary Fig. S1A). All members have the same number of chromosomes, a high degree of genomic synteny, and are to some degree crossable (Bedinger *et al*., 2011; Knapp and Peralta, 2016). Wild species naturally occur on the western slopes of the Andes in generally dry environments, and span a 3000 km North-South range (Supplementary Fig. S1B). The close relationship between species with red-orange fruits *S. lycopersicum, S. pimpinellifolium, S. galapagense* and *S. cheesmaniae* (“red-orange fruit clade”) has been robustly proven (Peralta *et al*., 2008 and Knapp and Peralta, 2016).

We have profiled CBC intermediates in five species from the *Solanum* sect. Lycopersicon (Supplementary Fig. S1A): *S. lycopersicum, S. pimpinellifolium, S. cheesmaniae, S. neorickii* and *S. pennelli*. The aims were to investigate if CBC intermediate profiles vary between closely-related species, to ask if the variance relates to phylogeny and to compare their profiles with those previously reported for phylogenetically-distinct species.

## Material and Methods

### Plant growth and harvest

*Solanum lycopersicum* (var. ‘Moneymaker’), *Solanum pimpinellifolium, Solanum cheesmaniae, Solanum neorickii* and *Solanum pennellii* seeds were provided by Saleh Alseek. Seeds were germinated on 2% v/v sucrose agar and grown in a 16h day/8h night (irradiance 590 µmol photons m^-2^ s^-1^, 23°C/20°C, 60% relative humidity). The apical leaf blade of the third fully-expanded leaf was harvested under ambient light by quenching in liquid nitrogen without shading. For each sample, apical leaves from two plants were randomly pooled prior to grinding and further processing. The number of replicate samples was S. *lycopersicum* n=6, *S. pimpinellifolium* n=6, *S. cheesmaniae* n=5, *S. neorickii* n =6, *S. pennellii* n=7. Each replicate was with leaves from different plants.

### Metabolite extraction and analysis

Frozen leaf material was homogenized with an oscillating ball mill (Retsch, Haan, Germany; https://www.retsch.com) and metabolites extracted and quantified by LC–MS/MS as in Arrivault *et al*. 2009; 2019). Stable-isotope labelled internal standards were added to correct for matrix effects (Arrivault *et al*., 2015). 3PGA was determined enzymatically in freshly prepared trichloroacetic acid extracts (Jelitto *et al*., 1992) using a spectrophotometer (Shimadzu, Kyoto, Japan; www.shimadzu.de).

### Statistical analysis

Statistical analysis (ANOVA and Tukey’s test) was performed in GraphPad Prism version 10.2.2. The z-score normalization (scaling the data for each metabolite to have a mean of 0 and a standard deviation of 1) for the PC plots was carried out in RStudio version 2025.09.2+418.

## Results

### Experimental design

CBC metabolites were profiled in the apical leaf blade of the third fully expanded leaf, harvested under growth irradiance. For each species, 5-7 replicate samples were harvested from different plants. Metabolite levels were initially expressed on a fresh weight (FW) basis. However, there can be systematic changes in the levels of all metabolites if leaves from different genotypes have differing water contents due, for example, to differing protein content and/or cytoplasmic volume and/or cell wall contribution. To exclude these secondary effects, in each sample the amount of C in a given metabolite was expressed as a percentage of the summed C in all measured metabolites (Supplementary Table S1). In previous publications (Arrivault *et al*., 2019) this normalization was termed a ‘dimensionless’ data set. Xu5P and Ru5P are not separated by the LC-MS/MS platform and are therefore represented together (Xu5P+Ru5P). DHAP, which is the major triose phosphate was detected, but not glyceraldehyde 3-P.

### Comparison of CBC metabolite levels in five different *Solanum* sect. Lycopersicon species

Fig. 1A shows the normalised metabolite levels in each genotype as boxplots. A Tukey post-hoc test was performed to detect significative inter-species differences. There were many significant inter-species differences in the 10 pairwise comparisons, especially for RuBP (5), FBP (5), S7P (4), R5P (4) and 2PG (3), with less for DHAP (2) and SBP (2) and Ru5P+Xu5P (1) and none for 3PGA and F6P. (Supplementary Table S2). Pairwise species comparisons pointed to *S. pennellii* being the most divergent species (in each comparison, 4-6 metabolites showed significant differences) followed by *S*.*pimpinellifolium* (1-6 metabolites) and S. *lycopersicum* and *cheesmaniae* (0-4 metabolites). No significant differences were found between metabolites when *S. lycopersicum* and S. *cheesemaniae* were compared (Supplementary Table S2).

**Figure 1.**
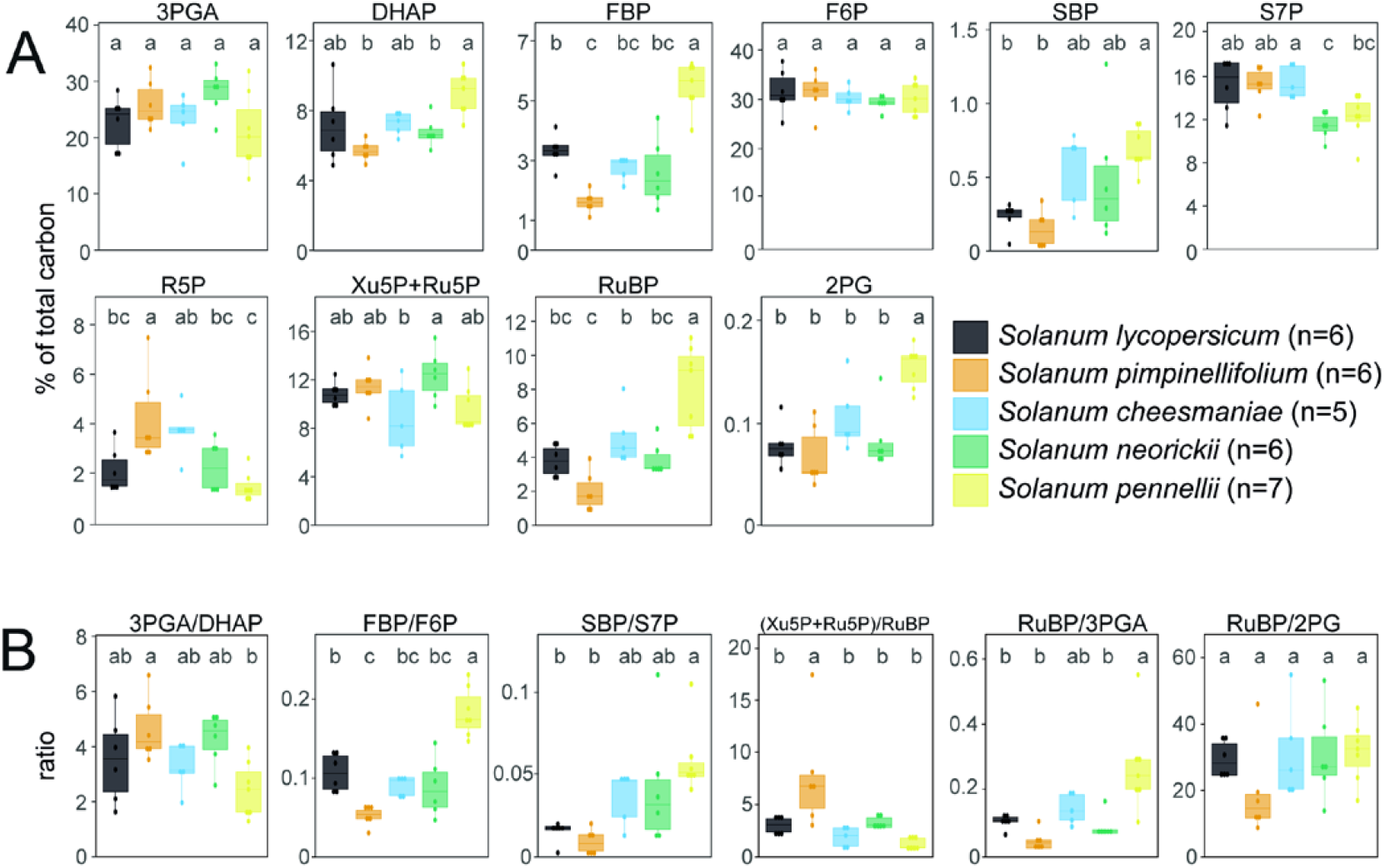
Levels of CBC intermediates and 2PG, and their ratios in five *Solanum* sect. *Lycopersicon* species. (**A**) Metabolite contents. (**B**) Metabolite ratios. The 3PGA/DHAP ratio is a proxy for the balance between energy supply from the light reactions and energy consumption in the CBC. The FBP/F6P, SBP/S7P and (Xu5P+Ru5P)/RuBP ratios are proxies for resistance to flux at FBPase, SBPase and PRK, respectively. The RuBP/3PGA and RuBP/2PG ratios indicate resistance at the carboxylation and oxygenation reactions of Rubisco. Metabolite levels were normalised by expressing the C in a given metabolite as a percentage of the summed C in all CBC metabolites in that species (termed a ‘dimensionless’ data set, see Arrivault *et al*., 2019). To do this, in a given sample, the level of each metabolite was first transformed to C equivalent values by multiplying the amount (nmol g FW^−1^) by the number of C atoms in the metabolite. The C equivalent amounts of all CBC intermediates plus 2PG were then summed for that sample. In the last step, the C equivalent value of a given metabolite was divided by the summed C equivalent value. Between 5 and 7 replicate samples were harvested and analysed for each species. Single datapoints are shown on the graphs, the upper and lower limits of the boxes represent the third and first quartile, and the whiskers represent the interquartile range. The letters on the graph indicate significant differences between the means of each species, with p-value < 0.05 (ANOVA and Tukey’s test performed in GraphPad Prism version 10.2.2, graphs generated in Rstudio version 2025.09.2+418). Note the different scales on the y-axes.

To provide an integrated overview, principal components (PC) analysis was performed on z-score normalized data. PC1, PC2 and PC3 accounted for most of the total variation (41.5%, 24.9% and 14.2%, respectively). The two major PC are shown in Fig. 2. PC analysis separated *S. pennellii* from the other species. *S. pimpinellifolium* was also quite separated from other species, and took a diametric position to *S. pennellii*. The other three species overlapped with each other. Separation was mainly along PC1, driven by opposed changes of RuBP, SBP, FBP, 2PG and DHAP, which aligned with *S. pennellii*, and R5P, Xu5P+Ru5P, S7P and 3PGA, which aligned with *S. pimpinellifolium*. There was no evident separation of species in PC2.

**Fig. 2.**
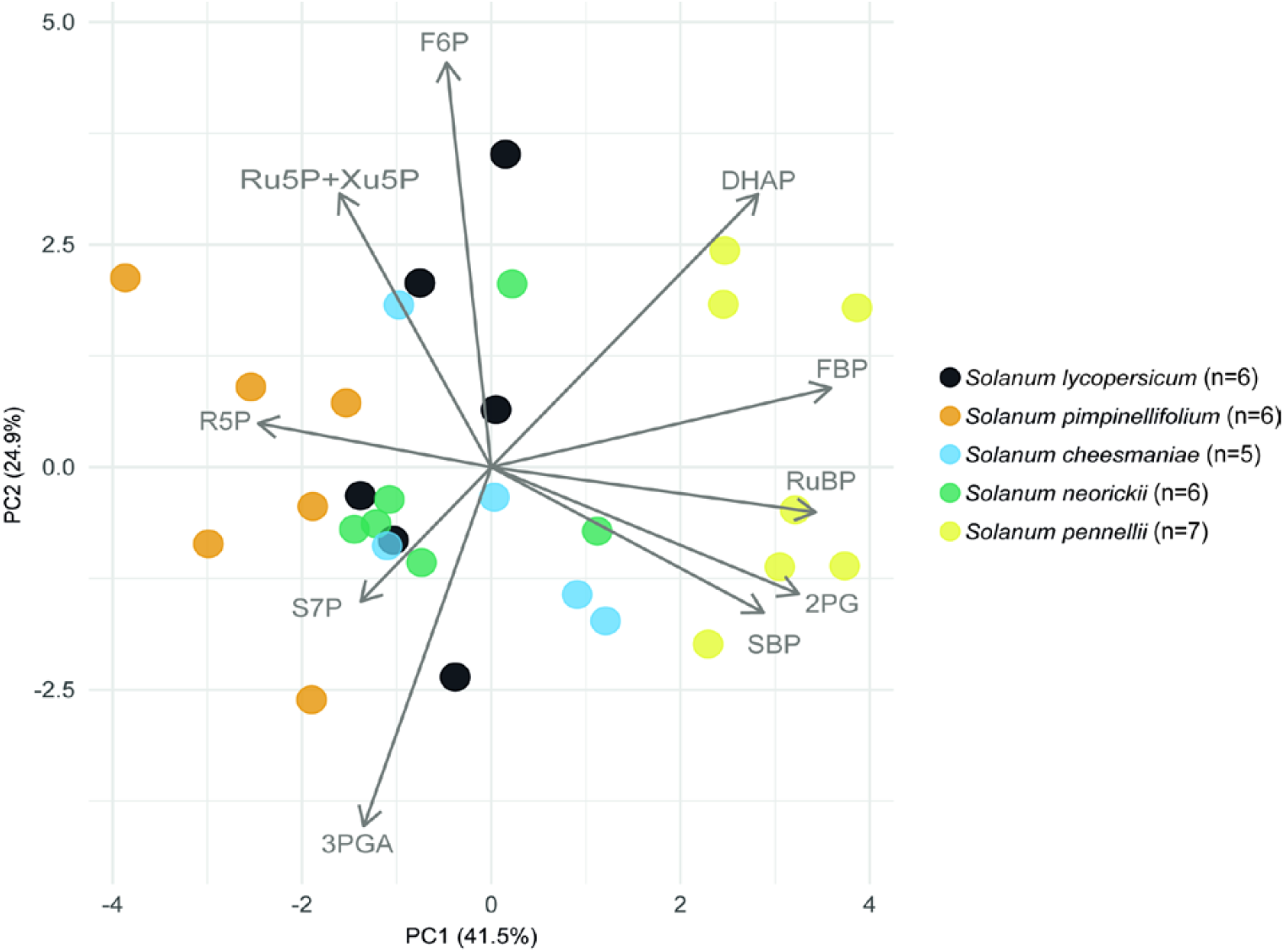
PC analysis of metabolite profiles from the five *Solanum* sect. *lycopersicon* species. Metabolite amounts were transformed in a dimensionless dataset (see legend of Fig. 1). Before the PC analysis, the values were z-score normalised, scaling the data for each metabolite to have a mean of 0 and a standard deviation of 1. The z-score normalization and graph generation were carried out in RStudio version 2025.09.2+418. The display shows each replicate sample, and the vectors for CBC metabolites that drive separation of the samples.

### Comparison of metabolite ratios in five different *Solanum* sect. Lycopersicon species

Metabolite ratios reflect relative abundance of the substrate and product of a reaction, and provide a proxy for restriction on flux. The 3PGA/DHAP ratio (Fig. 1B), which is indicative of energy supply from the photosynthetic light reactions, was largely conserved among species. The only significant pair-wise difference was between a slightly lower value in *S. pennellii* and a slightly higher ratio in *S. pimpinellifolium*. Neither of these species was significantly different to the other three species. The FBP/F6P, SBP/S7P, and (Ru5P+Xu5P)/RuBP ratios (Fig. 1B) provide information for the reactions catalysed by at FBPase, SBPase and PRK, respectively. The FBP/F6P ratio was significantly higher in *S. pennellii* and lower in *S. pimpinellifolium* than in the other species. The SBP/S7P ratio was also significantly different higher in *S. pennellii* than *S. pimpinellifolium*, and showed a similar but non-significant trend to the FBP/F6P ratio for the other three species. *S. pimpinellifolium* displayed a significantly higher (Ru5P+Xu5P)/RuBP ratio than the other four species. The RuBP/3PGA and RuBP/2PG ratios (Fig. 1B) reflect Rubisco carboxylation and oxygenation, respectively *S. pennellii* showed a significantly higher RuBP/3PGA ratio than S. *lycopersicum, S*.*pimpinellifolium* and *S. neorickii*, and a non-significant trend to a higher value than S. *cheesmaniae*. No significant differences were found for the RuBP/2PG ratio.

### Comparison of *Solanum* sect. Lycopersicon species with phylogenetically-distinct C_3_ species and C_4_ species

A second PC analysis was performed after combining the dataset for the five *Solanum* species with published datasets for five other C_3_ species (*A. thaliana, N. tabacum, O. sativa, T. aestivum, M. esculenta*) from Arrivault *et al*. (2019). This analysis again used z-score normalised data. Fig. 3A shows PC1 and PC2, which accounted for 47.2% and 22.7% of total variation, respectively. The *Solanum* species, in particular *S. pennellii* and *S. pimpinellifolium*, remained partly separated from each other in this combined analysis. All of the *Solanum* species were clearly separated from the other C_3_ species. The closest was *A. thaliana*. whilst *T. aestivum, O. sativa, N. tabacum* and *M. esculenta* were strongly separated from the *Solanum* species. The separation was seen mainly in PC1 and was driven by higher F6P, S7P, R5P and 3PGA to a certain extent DHAP and lower RuBP, SBP, FBP and to a certain extent 2PG and Xu5P+Ru5P in the Solanum species than the other C_3_ species. There was no consistent separation in PC2.

**Fig. 3.**
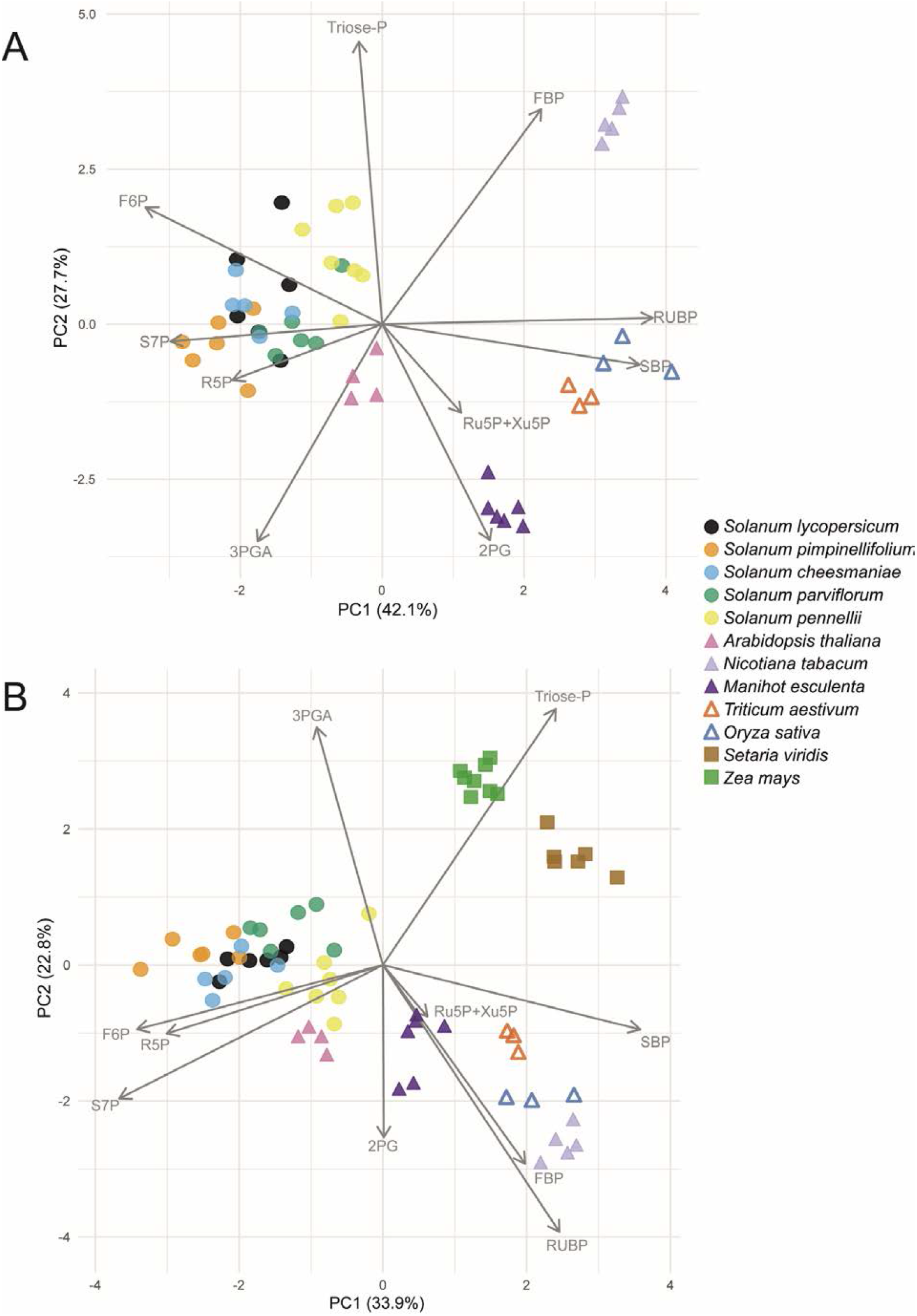
PC analysis of CBC metabolite profiles from *Solanum* sect. *lycopersicon* species together with further published data sets. (**A**) Together with five further C_3_ species and (**B**) together with these five C_3_ species and two C_4_ species. Metabolite amounts were transformed into a dimensionless dataset for each sample (see legend to Fig. 1). Before the PC analysis, the values were z-score normalised (see legend to Fig. 2). The displays show each replicate sample, and the vectors for CBC metabolites that drive separation of the samples.

A third PC analysis was performed including the five Solanum species, the four above C_3_ species and two C_4_ species (*Zea mays, Setaria viridis)* again using data from Arrivault *et al*. (2019). Fig. 3B shows PC1 and PC2, which accounted for 36.2% and 25.4% of total variation, respectively. As seen in earlier studies (Arrivault *et al*. 2019; Borghi *et al*., 2022), C_3_ species separated from C_4_ species, driven mainly by higher 3PGA and/or DHAP levels in the C_4_ species. Even in this analysis of species with diverse modes of photosynthesis, the *Solanum* species are separated from other C_3_ species although again lying quite close to *A. thaliana*, and show partial separation between themselves.

## Discussion

The tomato *Solanum* sect. Lycopersicon clade was used to test, firstly, if there is interspecies variation in CBC metabolite levels in a set of closely phylogenetically related species and, secondly, if the CBC metabolite profiles of this set of phylogenetically related species are distinct from or overlap with those of phylogenetically distant C_3_ species.

There were many significant differences in levels of individual CBC intermediates between the five Solanum species, especially for RuBP and FBP that showed five and S7P and R5P that showed four significant differences in a total of ten pair-wise comparisons (Fig. 1A, Supplementary Table S2). *S. pennelli* was the most divergent species, followed by *S. pimpinellifolium*, whilst *S. lycopersicum* ‘Moneymaker’ and *S. cheesmaniae* were the most similar. In a PC analysis of CBC metabolite profiles, *S. lycopersicum* ‘Moneymaker, *S. cheesmaniae*, and *S. neorickii* overlapped, and *S. pimpinellifolium* and especially *S. pennellii* grouped separately and were also at the opposing ends of the overall distribution (Fig. 2). These observations reveal variation in poising of the CBC between closely-related species. This complements other research that has identified variation in light acclimation, stomatal response, energy dissipation and further photosynthetic traits between closely related or even within species (Jakobson *et al*., 2016; van Rooijen *et al*., 2017; Flood, 2019; Kromdijk *et al*., 2022, Sakoda *et al*., 2022).

Separation of species in the PC analysis was mainly driven by opposed changes in RuBP, SBP, FBP, 2PG and DHAP (higher in *S. pennellii*) and R5P, Xu5P+Ru5P, S7P and 3PGA (higher in *S. pimpinellifolium)*. To investigate which reactions might be responsible, metabolite ratios were calculated (Fig. 1B). The 3PGA/DHAP ratio did not change significantly except for somewhat lower values in *S. pennellii* compared to *S. pimpinellifolium*, indicating that the balance between energy provision by the light chloroplast electron transport chain and energy consumption in the CBC has shifted in favour of the former in *S. pennellii*. Within the CBC, *S. pennelli* was characterised by higher substrate/product ratios (indicating higher resistance to flux) at FBPase, SBPase and Rubisco, whilst *S. pimpinellifolium* showed increased resistance to flux at PRK. The other species were intermediate between *S. pennellii* and *S. pimpinellifolium*. Lana-Costa *et al*. (2020) recently reported a QTL in an *S. pennelli* vs *S. lycopersicum* introgression line population that was associated with higher rates of photosynthesis and increased Rubisco activity. It is tempting to suggest that the and the higher FBP/F6P and SBP/S7P ratios and higher RuBP in *S. pennellii* in our study in might reflect increased resistance to flux at FBPase and SBPase in *S. pennellii* and, possibly, increased binding of RuBP at more abundant Rubisco active sites.

Our observations are broadly consistent with the hypothesis that CBC metabolite profiles differ due to lineage-dependent adaptation in closely-related C_3_ species, as the most divergent profiles were found in *S. pennelli*, which is more phylogenetically distant compared to the other species (Supplementary Fig. S1A). Within the group of more closely related species, the observed CBC metabolite profiles do not follow phylogeny so closely. In particular, the cultivated *S. lycopersicum* (variety Moneymaker) is very similar to *S. cheesmaniae* and *S. neorickii* but distinct from its progenitor *S. pimpinellifolium. S. pimpinellifolium* spans a long North-South axis along the Andes and has also spread from wet climate zones in coastal Ecuador to much drier zones in central and southern Peru (Lin *et al*., 2020, Supplementary Fig. S1B).

The *S. pimpinellifolium* accession used in this study (LA1589) has been described as endemic to centre-South area of Peru (Lin *et al*., 2020), while it is thought that the modern tomato derived from genotypes from more northerly populations (Lin *et al*., 2020), possibly explaining the metabolic space between *S. pimpinellifolium* and *S. lycopersicum*. Profiling of additional *S. pimpinellifolium* and *S. lycopersicum* varieties could ascertain whether the differences in CBC poise found in this study are due to genetic diversity of the starting breeding material or a consequence of the breeding process itself, as it is possible that linkage drag with traits that ‘were bred for resulted in a different CBC poise…Furthermore, it is possible that the variance in CBC profiles uncovered in this study originated from the differing ecological niches that the Solanums occupy, having adapted to different (micro) climates or nutrient supply (Flood, 2019). They might be explained by evolution after the first speciation event, possibly including adaptation to different climates (Peralta *et al*., 2008; Pease *et al*., 2016; Gibson and Moyle, 2020; Lin *et al*., 2020). Adaptation to microclimate as a driver of (micro-)evolution has been discussed for *A. thaliana* (Hancock *et al*., 2011; Kubota *et al*., 2015). However, it cannot be excluded that differences originated due to random genetic drift (Lynch *et al*., 2016) in common progenitors and were then exacerbated by geographical separation of populations (Bock *et al*., 2023). Metabolic profiling of another set of closely related C_3_ species could reveal if they also exhibit diversity in CBC poising, or whether it is a characteristic of the *Solanum* clade.

When the tomato clade is compared to other C_3_ species, their CBC metabolite profiles clustered together, separately from non-Solanum species. At the same time, variation can be seen in how the other four C_3_ species distribute. This probably points out to a certain degree of conservation of CBC poise in Solanum species, which jointly share a different evolutionary trajectory to the other investigated C_3_ species. The dicot *A. thaliana* was most similar to the Solanum species, whilst the dicot *N. tabacum* and *M. esculenta* and the monocot C_3_ species *O. sativa and T. aestivum* were much further separated from the *Solanum* sect. Lycopersicon species. When the experimental space is expanded by including C_4_ species the Solanum species still group separately from other C_3_ species. Overall, these observations are consistent with the idea that the CBC diverged in a phylogenetically-related manner. However, *N. tabacum* grouped further away than *A. thaliana*, even though it is a member of the Solanaceae family and thus more closely phylogenetically-related to *Solanum* sect. Lycopersicon, indicates that the extent of separation of CBC metabolite profiles is not solely due to phylogeny.

In conclusion, variation in CBC metabolites profiles within *Solanum*, together with their separation from other C_3_ and C_4_ species, indicates that CBC evolution is shaped both by phylogenetic relatedness and lineage-specific adaptation. Species-variation in the Solanum might be further investigated using introgression lines of wild *Solanum* species (de Oliveira Silva *et al*., 2018; Lana-Costa *et al*., 2020). More generally, there is much interest in genetically improving photosynthesis (Long *et al*., 2025). Species-variation in poising of the CBC might be taken into account when designing strategies to improve photosynthesis in crops.

## Abbreviations

2PG: 2-phosphoglycolate
3PGA: 3-phosphoglycerate
CBC: Calvin-Benson cycle
CO_2_: carbon dioxide
DHAP: dihydroxyacetone phosphate
F6P: fructose 6-phosphate
FBP: fructose 1,6-bisphosphate
FBPase: fructose-1,6-bisphosphatase
LC-MS/MS: liquid chromatography coupled with tandem mass spectrometry
Liquid N_2_: liquid nitrogen
O_2_: molecular oxygen
PC: principal component
PRK: phosphoribulokinase
R5P: ribose 5-phosphate
Ru5P: ribulose 5-phosphate
Rubisco: ribulose-bisphosphate carboxylase-oxygenase
RuBP: ribulose 1,5-bisphosphate
S7P: sedoheptulose 7-phosphate
SBP: sedoheptulose 1,7-bisphosphate
SBPase: sedoheptulose-1,7-bisphosphatase
Xu5P: xylulose 5-phosphate

## Supplementary material

**Supplementary Fig. S1**. Tomato clade (*Solanum* sect. *lycopersicum*) phylogenetic tree and geographical distribution of wild species

**Supplementary Table S1**. Content of CBC intermediates and 2PG in species of *Solanum* sect. *Lycopersicon*.

**Supplementary Table S2**. Pairwise species comparison of metabolite levels with Tukey post-hoc test p-values

## Acknowledgment

The authors thank Dr. Saleh Alseek from the Max-Planck Institute of Molecular Plant Physiology for providing the *Solanum* spp. seeds.

## Data availability

All data in this study are presented in the Supplementary material which is available online.

## Author contribution

VC and MS conceived the experiments. VC grew plants, harvested samples and extracted them for downstream analysis. VC and SA processed samples by LC-MS/MS and analysed the results. VC performed statistical analysis and generated all graphs. VC wrote the original draft and all authors reviewed and edited it. MS supervised the project.

